# Where Climate Change and Sampling Bias Collide: Challenges of Predicting Biodiversity Change in Canada

**DOI:** 10.64898/2025.12.11.693756

**Authors:** Noah Wightman, Isaac Eckert, Brian Leung, Laura J. Pollock

**Affiliations:** Department of Biology, McGill University; Quantitative Life Sciences, McGill University

## Abstract

Anticipating biodiversity change is critical in rapidly warming regions, yet challenging because these areas often coincide with poor sampling. Data gaps are widely understood to interfere with species distribution models (SDMs), but this is difficult to detect with biased data. We test SDM bias-correction methods with a new occurrence-checklist-range (OCR) validation approach and evaluate prediction discrepancy for ∼700 Canadian terrestrial vertebrate species. We found: 1) bias-correction improves model performance against independent (checklist and range) data, but not against typical occurrence cross-validation, 2) predicted richness differed among methods (up to 2.7-fold), especially in the north, and 3) counterintuitively, future projections varied less (by 28%) because well-sampled climate space will shift north. Our findings suggest potential widespread overconfidence in SDM predictions for the unevenly sampled world, with implications for the growing reliance on biodiversity estimates for planning and policy. OCR validation and methodological discrepancy measurements are relatively easy ways to address this.

## Introduction

Projections of biodiversity shifts are critical yet challenging for regions experiencing rapid climate change. These regions provide invaluable ecosystem services such as timber, freshwater, and carbon sequestration, but future predictions suffer from large uncertainty arising from climate models, novel climatic conditions, dispersal abilities, and changing biotic interactions (Brodie *et al*. 2022; Thuiller *et al*. 2008). Compounding the problem, many of the regions anticipating the most severe climate change also have the poorest biodiversity sampling (e.g. rapid warming in the arctic and boreal forest, precipitation increase in central Africa, soil drying in the Amazon; IPCC. 2023). This unfortunate pairing of rapid climate change and data limitation likely has under-explored consequences on the SDM predictions used to inform conservation assessments and policy (Brodie *et al*. 2022, KMGBF 2022).

Sampling bias is a known challenge for SDMs. SDMs estimate species’ environmental preferences by relating species occurrence data to environmental covariates and projecting these relationships in geographic space to estimate the distribution of suitable habitat (Elith & Leathwick 2009; Franklin 2010). However, occurrence data are often biased towards accessible locations, such as cities, roads, and nature reserves (Beck *et al*. 2014). Models can mistakenly assume that species prefer these well sampled environments (Baker *et al*. 2022), causing predicted distributions to mirror that of sampling effort rather than the true range of the species (Kramer-Schadt *et al*. 2013; Syfert *et al*. 2013). This bias is particularly problematic for the many species with mainly presence-only observations (from citizen science, museum collections, government agency atlases, etc.), rather than systematic presence-absence surveys (Phillips *et al*. 2009). Many methods have been developed to correct for sampling bias, including filtering occurrence data (Hidalgo-Mihart *et al*. 2004), balancing it with biased background data (Phillips *et al*. 2009), or explicitly modelling and factoring out the bias (Leung *et al*. 2019; Warton *et al*. 2013). In principle, these methods improve distribution estimates, but we do not fully understand the limitations to their performance (i.e. (Liu *et al*. 2020; Mendes *et al*. 2020), how to evaluate these, and whether predictions differ among bias correction approaches.

In general, theoretical and empirical studies show that common bias-correction methods can improve SDM predictions and produce reliable predictions (Barber *et al*. 2022; Inman *et al*. 2021; Kramer-Schadt *et al*. 2013; but see Baker *et al*. 2024). But simulation studies comparing bias-correction methods typically either simulate a smooth gradient of sampling effort (ie Baker *et al*. 2022) or mimic the sampling of relatively well-sampled regions like Europe and USA (i.e. Dubos *et al*. 2021; Varela *et al*. 2014). Real-world comparison studies of SDMs have also focused on these well-sampled regions (Barber *et al*. 2022; Fourcade *et al*. 2014) when globally, there are large ecologically important areas that have much thinner sampling coverage (such as the arctic and boreal forest, most of Africa, the Amazon basin, etc.; Meyer *et al*. 2015). We need to better test the consequences of large, but realistic data gaps on SDM performance.

A major challenge is that data limitations make it difficult to properly evaluate *model performance.* Typical cross-validation approaches use hold-out sets from the original occurrence data, and thus will include the same biases, leading to skewed results even in the best-sampled conditions (Roberts *et al*. 2017; Syfert *et al*. 2013). Validating predictions with fully independent datasets is considered the gold standard but rarely done in practice as independent point-based datasets are uncommon. However, other types of datasets can be used for evaluation at broad scales (range-level model performance) by comparing them to expert-derived range maps, including expert review (Boyd *et al*. 2023) or globally available IUCN ranges (Fourcade 2016). Alternatively, full species checklists from expert knowledge of a particular location (such as protected areas) represent presence-absence data that are often available in remote areas (Edwards *et al*. 1996), and could evaluate finer-scale patterns within a range (checklist-level model performance). Here, we use a multifaceted, occurrence-checklist-range (OCR) approach to validation that could prove especially valuable in data-sparse regions.

Even models that each validate well can produce very different predictions, so the discrepancy between different methods (model uncertainty) is also important to consider. Predictions are often used to inform conservation decisions that account for future changes (Guisan *et al*. 2013). However, predictions that are sensitive to methodological choices of bias correction must be interpreted with caution. To make decisions effectively, we need to understand where and when differences among methods (which we refer to as *prediction discrepancy)* are most pronounced (Grimmett *et al*. 2020).

Here, to understand the real-world consequences of highly biased data on SDM predictions, we comprehensively test how bias-correction methodology affects species distribution estimates and range shift projections for over 700 terrestrial vertebrates of Canada - a large country with boreal and arctic regions that are expected to warm at a rate 3 times faster than the global average (Bush & Lemmen 2019) and sparse, extremely biased biodiversity data (Hughes *et al*. 2021). We evaluate two aspects of model quality—*model performance* and *prediction discrepancy*. Specifically, we: 1) evaluate *model performance* of different bias-correction methods using both occurrence cross validation and independent datasets, and 2) assess how *prediction discrepancy* varies across space and time by comparing methods’ predictions of current and future biodiversity. If we assume based on previous findings (Barber *et al*. 2022; Inman *et al*. 2021) that predictions from bias-correction methods are reliable, then we expect that bias-correction methods should perform better than with no correction and result in little discrepancy in predictions. We also expect that SDM predictions should differ *more* in the poorly sampled north compared to well-sampled south and differ *less* in the present compared to future projections.

## Methods

### Occurrence data

We focused on terrestrial vertebrate species in Canada because they reflect ecosystem health (Loyola *et al*. 2007), are a focus of conservation efforts (Roberge & Angelstam 2004), and have relatively complete data (Troudet *et al*. 2017). We obtained species occurrences from the global biodiversity information facility (GBIF. 2022), which compiles presence-only data from many sources. We curated a species list of all terrestrial vertebrates recorded on GBIF in Canada, and removed all non-native, extinct and domestic species, yielding 718 study species (466 birds, 155 mammals, and 80 herptiles). We extracted occurrence data from 1970 to present for mammals and herptiles, and from 1990 to present for birds. This prioritized recent data with high geographic precision for the well-sampled birds, and large enough sample sizes for the lesser sampled non-birds. For birds, we aimed to capture breeding ranges only, so we filtered occurrences to breeding season (June to July). If time-stamped data for a species was lacking (15% of birds), we instead filtered occurrences using Birds of the World breeding ranges (Cornell Lab of Ornithology, 2022). For all species, we further filtered out points with “urban” land cover (potential zoos/sanctuaries observations), and for 11% of species removed some visually obvious outlier points that were well outside the known range based on IUCN (2022) or expert knowledge, and were isolated from other points. To better sample Canada’s future climatic conditions, we sampled occurrences from both Canada and U.S.A. We gridded the occurrence data at a 1km^2^ resolution to match the climate data (each cell with at least one record was considered one occurrence). This geographic thinning is a preliminary way of reducing spatial bias. For species with >2000 occurrences, we sampled 2000 random occurrences for computational practicality.

### Environmental data

The environmental predictor variables included 5 climate variables, 2 topographic variables, and landcover. We used the following current bioclimatic variables from AdaptWest (2021); Mean Annual Precipitation, Degree-days below 0℃, Precipitation as snow, Hargreaves climatic moisture index, and Degree days above 18℃. These were selected for low collinearity and biological relevance. Current layers represent average climate from 1991-2020 from PRISM and WorldClim. Future climate is based on an ensemble projection of 13 climate models using the “business as usual” representative concentration pathway (RCP) 8.5 (the most severe changes to highlight the potential for prediction discrepancies). We also used topographic ruggedness index (from AdaptWest), topographic wetness index (derived from a 1km elevation map using “dynatopmodel” R package; Metcalfe *et al*. 2015), and a MODIS-based landcover layer reclassified to the following categories; unvegetated, hardwood forests, evergreen forest, mixed forests, shrubs, and grasslands (Friedl *et al*. 2010). Occurrence and environment data processing was the same as in Eckert *et al*. (2023).

To quantify the sampling coverage of environment space, we calculated the overlap between the distribution of sampled points and the distribution of overall landscape points for both Canada and USA. We used two important and intuitive dimensions of environment space (mean annual temperature and mean annual precipitation) for computational tractability and so that results could be visualized and easily interpreted. We projected 5km grids of Canada and USA into our 2D environment space, then obtained a smoothed distribution using kernel density estimation (kde2d function in “MASS” package in R; Venables & Ripley 2002). We then took the 95% confidence interval of this distribution (i.e. discarded the 5% of the volume with the lowest density) to obtain a 2D polygon representing each country’s environment space. We did the same with all sampled locations (each point was weighted by number of species sampled). Then, for each country, we calculated the proportion of its environment space polygon that was covered by the sampled environment space polygon. We then repeated this procedure using the future (2080, RCP 8.5) climate projections of Canada and USA (with the current sampling distribution to assess our data’s coverage of future conditions).

### Sample Bias Correction

We focused on 3 common approaches to correcting sampling bias; (1) Bias-covariate correction (BCC), (2) Target-group background (TGB), and (3) weighted target group background (WTGB). As a reference, we also fit SDMs with no correction (NC), using the presence data described above and a random sample of 10000 background points (each non-presence grid cell in the study area an equal chance of being selected).

We applied bias-covariate correction (BCC) by adding a “bias covariate” to each SDM (El-Gabbas & Dormann 2018; Warton *et al*. 2013). The covariate we used was a 30km sliding window average for density of GBIF points calculated based on occurrences of the entire species list. The bias covariate allows models to partition variation caused by sampling bias vs environmental preference. We then predicted corrected species distributions by setting the bias covariate to the maximum value everywhere. This removed the variance attributed to the bias covariate, so predictions should theoretically only be based on environmental preference.

The remaining two correction methods we used were target group background sampling (TGB) and weighted target group background sampling (WTGB). In these methods, the background sample was not random, but instead manipulated to have a similar bias as the presences. To do this, we selected background points only from grid cells where any species in a taxonomic “target group” has been recorded (we used all the species we modelled as a target group). In TGB, the background points had an equal chance of being sampled (without replacement) from any grid cell where at least one vertebrate had been recorded (Phillips *et al*. 2009). In WTGB, sampling probability was proportional to the total number of vertebrate occurrences recorded in that grid cell (Cerasoli *et al*. 2017). Thus, in highly sampled regions, there are also more absences, so the bias is “cancelled out” in the model fitting. Note that for all these methods (like all presence-only models), predictions are only *relative* estimates of probability of occurrence.

### Model Algorithms

For each species, we constructed 8 different SDMs (4 different bias correction methods each applied to 2 different model frameworks). Each bias correction method (NC, BCC, TGB, and WTGB) was used within both a boosted regression tree (BRT) model, and a Maxent model (BRT settings: tree complexity=4, learning rate=0.08, bag fraction=0.6, step size=200, Maxent settings: dismo default settings, but only allowed hinge features to limit flexibility). For comparability, species for which one or more of the models failed to converge were excluded from further analyses (698 species were kept). Maxent is the most widely used modelling framework for SDMs, and was designed specifically for presence-only data (Elith *et al*. 2010b; Phillips *et al*. 2006), while BRTs are becoming more popular, are flexible, and can resolve species preferences at fine scales (Elith *et al*. 2008). Using both model types allows us to draw conclusions about the consequences of each bias correction method independent of the modelling framework.

**Table 1:**
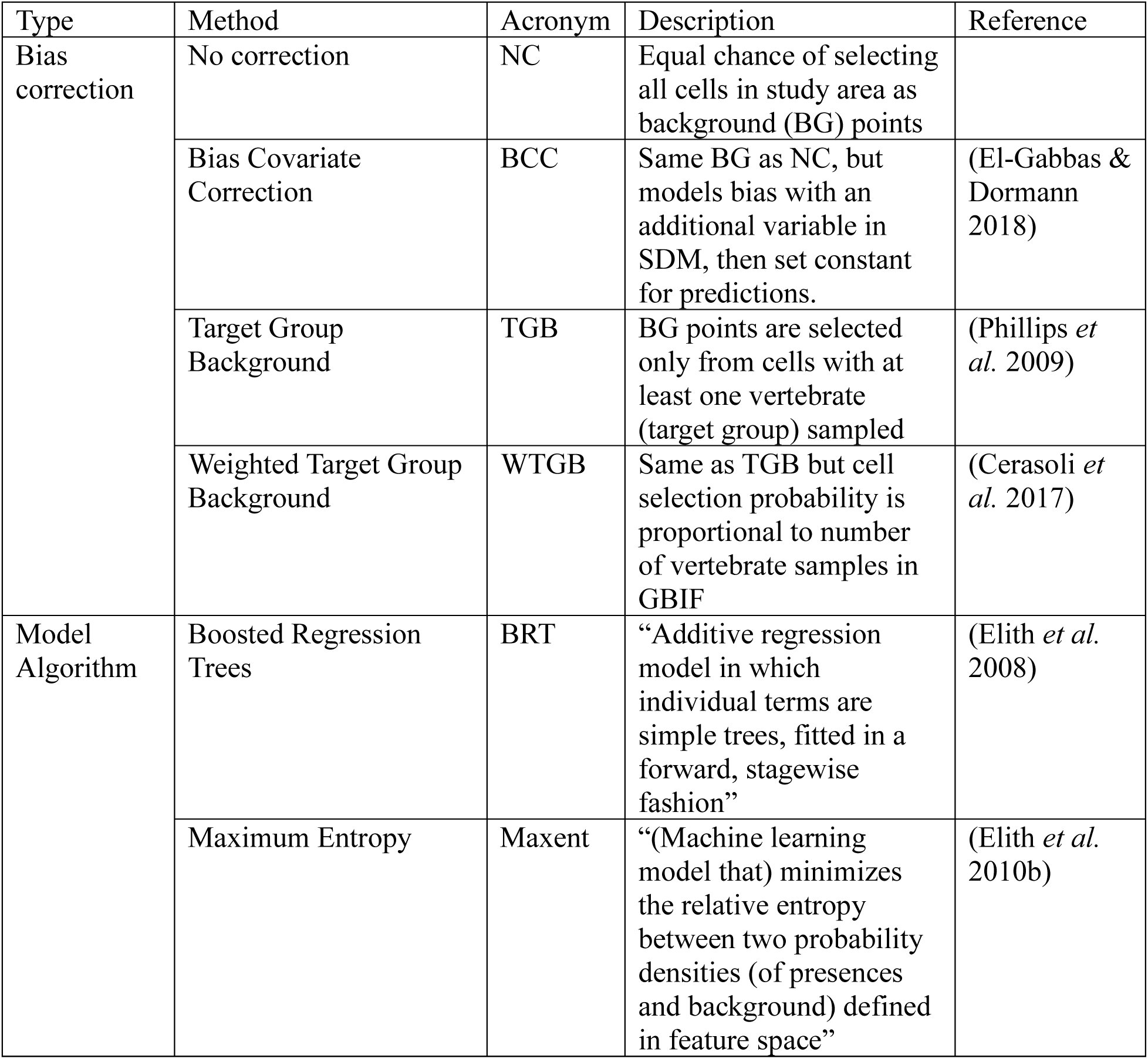
Summary of the sampling bias correction methods and modelling algorithms used to construct species distribution models in this study.

### OCR Model Validation

We validated models using a multifaceted approach which we call “occurrence-checklist-range” (OCR). All predictions and data were confined to Canada. We first used cross-validation using GBIF *occurrence* data. We trained models on 70% of the data and tested with the remaining 30% using Area under the receiver operator characteristic curve (AUROC), True Skill statistic (TSS) and Area under the precision-recall curve (AUPRC). We used models trained with 100% of the GBIF data for the remaining validations based on other independent data. As an independent site-level validation, we used (presence-absence) species *checklists* of national parks. We used the maximum predicted probabilities within each park and compared this to the observed presence/absences using AUROC, TSS and AUPRC. For birds, presences outside the expert breeding range were converted to absences (as we only modelled breeding range). Finally, to evaluate the models’ ability to capture *range* edges, we quantified the similarity between the IUCN range maps and predicted distributions using a minimum energy test (MET). This metric uses points sampled from two distributions and compares the within-distribution distances between points to the between-distribution distances (should be similar for similar distributions). Due to data limitations, 692, 548, and 541 species were successfully validated using held-out occurrence data, independent checklist data and range data respectively.

To simplify interpretation, we also calculated a combined validation score for each validation dataset. Although species-specific factors may affect predictive ability, we were interested in the relative performance of different methods *within* each species. Therefore, for the GBIF and Parks datasets, we centered each metric within each species (i.e., the 8 different models’ AUROC scores for a given species now have mean=0). We then normalized AUROC, TSS and AUPRC scores across all species (i.e. all AUROC scores now have mean=0 and s.d.=1) so that each metric is comparable. Finally, since all 3 metrics showed similar patterns (Fig. S1) we averaged the 3 metrics to get a combined validation score for each species and for each validation. The IUCN range dataset only used MET, which was normalized within each species. We also used a linear model to evaluate statistical differences between the 8 models’ validation scores. For each validation metric, we fit a model to the scores of all species with all 8 methods using validation score as the response variable, and bias correction method and modelling algorithm as categorical predictors, allowing us to separate the effects of corrections and algorithms.

### Prediction Discrepancy

To examine how sampling bias and climate change interact to influence model uncertainty (discrepancy between predictions), we used our 8 models to predict present and future (2080 with RCP8.5) species distributions at 5km^2^ resolution. All predictions were limited only to Canada (though U.S.A. was included in the model fitting to maximize sampled environment). To compare predictions among methodologies, we calculated expected current and future species richness for each method by summing (within each grid cell) the occurrence probabilities of all species. To measure discrepancy between methods, we calculated the coefficient of variation (variance/mean) of the species richness estimates (within each grid cell) across the 8 different models (for both current and future estimates).

We also examined how methodological choice affected range shift predictions. We used the current and future distribution predictions to estimate range expansion or contraction (future area/current area), and averaged (with a geometric mean because the metric is a proportion) across species to compare the overall differences between methods. We also calculated the percentage of the original range that remains stable in the future. For example, if a 1 km2 grid cell has a P(occupied) = 0.25 in the present and P(occupied) = 0.75 in the future, then it contributes min(0.25,0.75) = 0.25 km2 to the stable area.

## Results

### Sampling Bias in Environment Space

We compared Canada to the USA as an example of the relatively well-sampled conditions used in most SDM research. Sampling was notably more limited in environment space in Canada (18% of environmental space covered by sampling) compared to the United States (62% covered: Fig 1). Examining just the temperature and precipitation axes of environment space, sampling was mostly confined to the warmest, wettest corner of Canada’s environment space (Fig 1). Meanwhile, sampling covered the space well in the United States, although drier environments tended to be relatively under-sampled (Fig 1). This reflects geographic sampling patterns (sparse in cold northern Canada, somewhat sparse in deserts of western USA). In the future environment space, Canada increased to 29% of its projected environment space overlapped by sampling, as the well-sampled climates of the USA shift into Canada (Fig 1).

**Figure 1.**
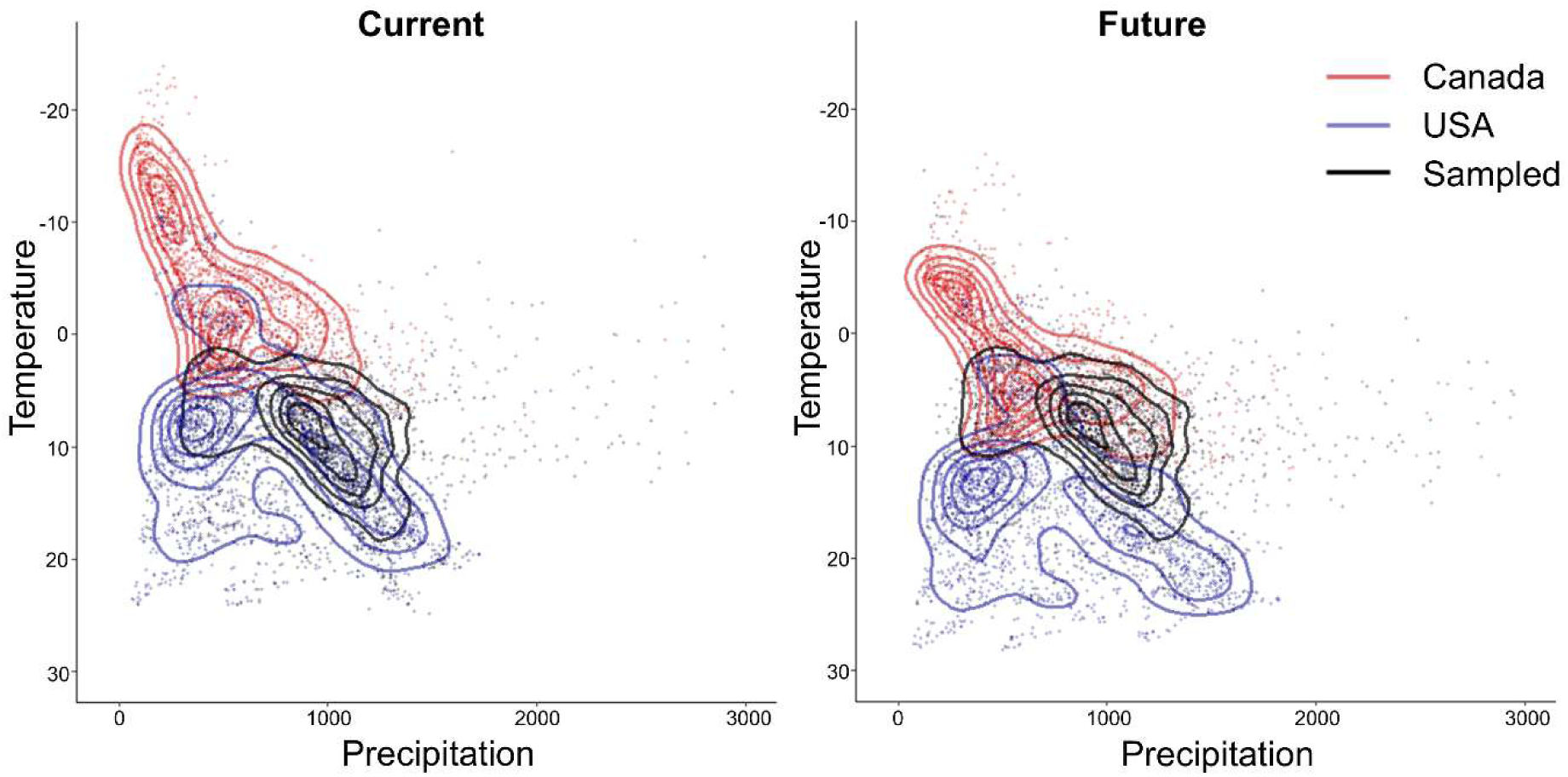
Sampling coverage of 2 dimensions of environment space (mean annual temperature and mean annual precipitation) for Canada and USA under current and future (2080, RCP 8.5) climate conditions. Black points represent 1km grid cells that have GBIF occurrences (weighted by number of species in the cell), while red and blue points represent randomly selected cells in Canada and USA respectively. Topographic lines represent a smoothed estimation of the 2D distribution of each set of points. In the “Future” panel, the sampled (black) points still represent the current climate of sampled grid cells.

### OCR Model Performance

To assess how different bias correction methods affect model performance, we compared the models against multiple datasets with an “Occurrence-Checklist-Range” validation. In cross-validation with occurrence data, bias correction methods performed worse than using no correction, with target group background (TGB) and weighted TGB (WTGB) performing worse than bias covariate correction (BCC) (Fig 2, Table 2). In contrast, using independent validation (checklist and range), bias correction methods performed better than uncorrected models, with BCC performing the best in both cases (Fig 2, Table 2).

**Figure 2.**
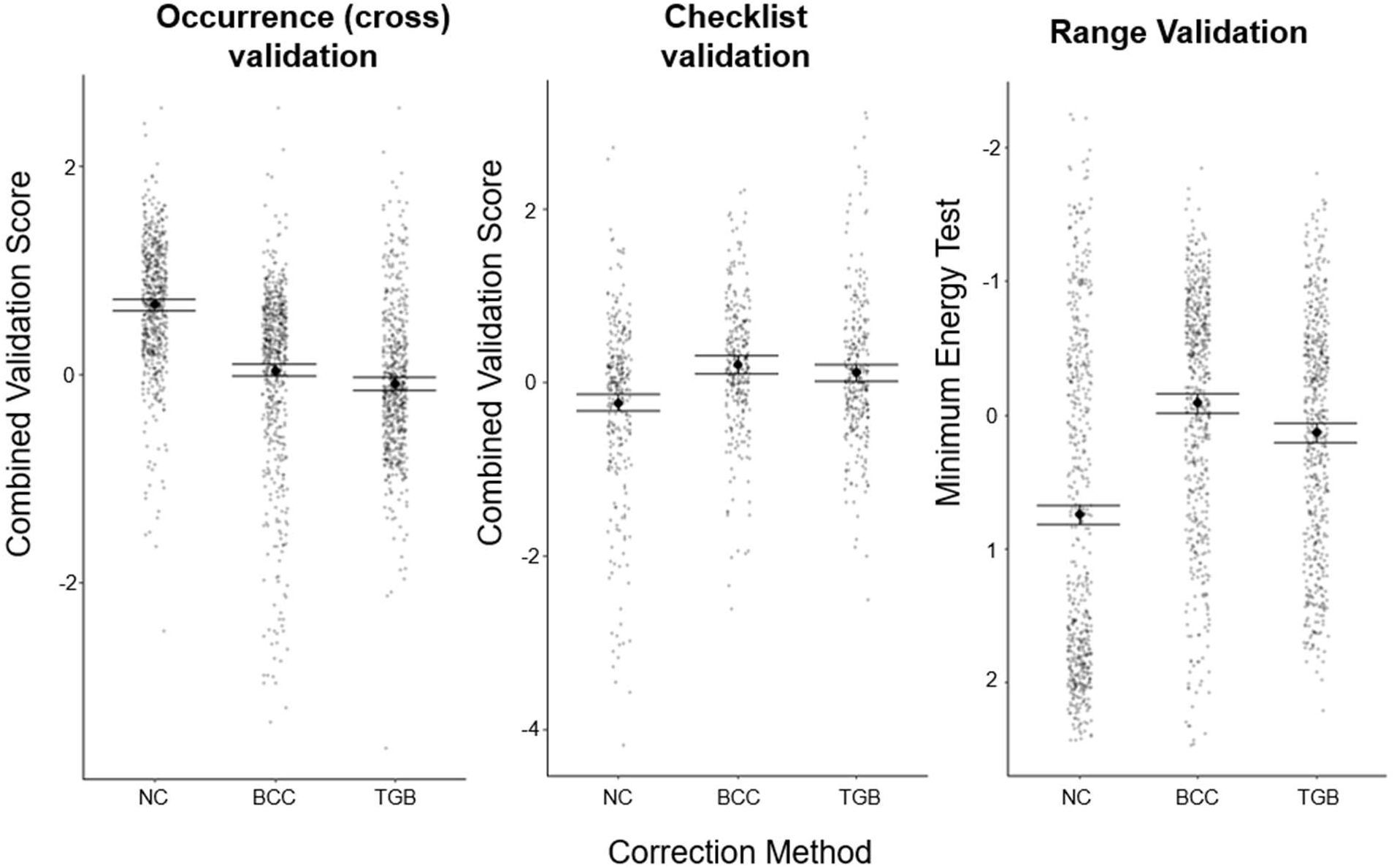
Performance of different bias correction methods (No Correction, Bias Covariate Correction, and Target Group Background) for 3 validation data sets (30% hold-out of the occurrences with pseudoabsences; cross-validation), presence/absence data from national park checklists, and range maps. Combined validation scores are averaged normalized AUROC, AUPRC and TSS metrics. Large black dots represent the mean for each method, with 95% confidence interval as bars. Note that the y-axis is reversed for range validation as lower values are better for this metric.

**Table 2:** Effects of bias correction methods and modelling algorithms on validation scores of species distribution models using multiple validation datasets. Datasets include 30% hold-out sets from the Global Biodiversity Information Facility occurrences with pseudoabsences (cross-validation), presence/absence data from Parks Canada national park checklists, and Range Maps from the International Union for the Conservation of Nature (or Birds of the World for birds). Combined validation scores are averaged normalized AUROC, AUPRC and TSS metrics. The intercept categories are No Correction for bias-correction and Boosted Regression Tree for Algorithm. Other bias-corrections are Bias covariate correction (BCC), Target group background (TGB) and Weighted TGB. Note that for IUCN only, negative effects mean model improvement.

|  | **Variable** | **Beta** | **SE** | **t-value** |
| --- | --- | --- | --- | --- |
| **OCCURENCE**<br>Combined<br>validation score | Intercept | 0.64 | 0.024 | 26.6 |
|  | Bias_cor BCC | -0.42 | 0.030 | -13.7 |
|  | Bias_cor TGB | -0.81 | 0.030 | -26.6 |
|  | Bias_cor WTGB | -0.86 | 0.030 | -28.2 |
|  | Algorithm Maxent | -0.24 | 0.022 | -11.1 |
| **CHECKLIST**<br>Combined<br>validation score | Intercept | -0.31 | 0.039 | -7.7 |
|  | Bias_cor BCC | 0.55 | 0.050 | 11.1 |
|  | Bias_cor TGB | 0.44 | 0.050 | 8.7 |
|  | Bias_cor WTGB | 0.45 | 0.050 | 9.1 |
|  | Algorithm Maxent | -0.11 | 0.035 | -3.1 |
| **RANGE**<br>Minimum<br>Energy Test | Intercept | 0.71 | 0.029 | 24.3 |
|  | Bias_cor BCC | -0.74 | 0.037 | -20.0 |
|  | Bias_cor TGB | -0.58 | 0.037 | -15.7 |
|  | Bias_cor WTGB | -0.54 | 0.037 | -14.7 |
|  | Algorithm Maxent | -0.49 | 0.026 | -18.8 |

BRT models outperformed maxent in occurrence (cross) and checklist validation but under-performed on ranges. However, both model algorithms showed similar patterns among bias correction methods for all validation datasets. Given the relative similarities between maxent and BRT, and with WTGB and TGB, we only report NC_BRT, BCC_BRT and TGB_BRT models (hereafter referred to as NC, BCC and TGB) in the following results

### Prediction Discrepancies

Beyond differences in validation, bias correction methods were also inconsistent when comparing predictions made by the models. Bias correction strongly altered predictions relative to uncorrected models and different bias correction methods varied strongly amongst themselves. We present discrepancies in predictions of 1) a species-specific distribution, 2) overall species richness, and 3) species-level climate driven range shifts.

#### Case Study: Ursus americanus

As an example, we look in detail at the distribution predictions of a wide-ranging large mammal in Canada, the Black bear (*Ursus americanus*). The NC model predicts that suitable bear habitat is mainly in southern Canada (even in the west it was generally hundreds of kilometers south of the IUCN-delimited northern range edge: Fig 3). In contrast, BCC predicted suitable habitat much further north, better matching the IUCN range, but also predicted moderate suitability well beyond the range edge (Fig. 3). Finally, TBG was intermediate, with suitable habitat throughout much of the range, but still declining well short of the northern range edge (especially in east). TGB also had more patches of unsuitable habitat elsewhere within the range, and very little suitable habitat beyond the range edge (Fig 3).

**Figure 3.**
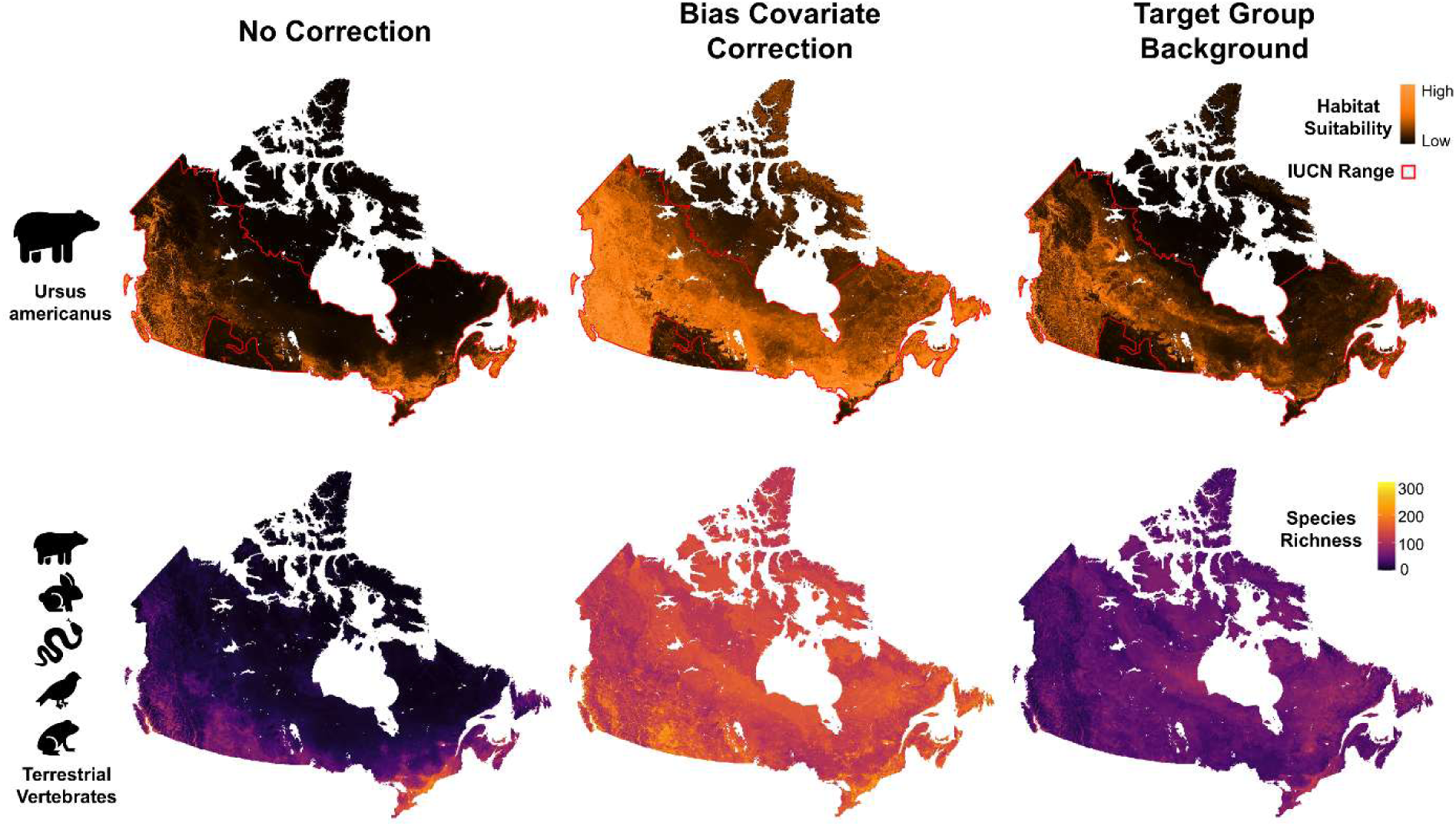
Differences between three bias correction methods for predictions of Boosted regression tree species distribution models (SDMs) across Canada. Top row: Predicted habitat suitability for an example species, the Black bear (*Ursus americanus*), with its range (delimited by the international Union for the conservation of Nature) overlaid as a realism check for SDM predictions. Bottom row: Estimated current species richness (as summed single species SDM predictions) of 698 terrestrial vertebrate species native to Canada.

#### Species Richness

NC models predicted high species richness in densely inhabited areas of southern Canada, and extremely low species richness in remote northern regions (Fig 3). TGB models showed a less concentrated pattern; they still tended to predict high richness near cities in the south, but in remote regions they predicted higher richness than NC (including high richness near the treeline: Fig 3). In contrast, BCC models predicted high species richness fairly evenly throughout all of Canada (Fig 3). Projections of future richness largely represented northward shifts of the current richness patterns (Fig. S2).

Predictions of mean species richness across the landscape differed correspondingly; NC models predicted only 17.7 (s.d. 26.0) species per grid cell, while TGB and BCC predicted species richness at 46.3 (s.d. 19.4) and 126.8 (s.d. 37.2) respectively. Change in mean richness between present and future varied also among methods. NC predicted richness to increase in the future to 34.4 (s.d. 32.2, 94% increase from present), while TGB and BCC predicted decreases in species richness to 37.2 (s.d. 21.1, 20% decrease) and 70.8 (s.d. 23.4, 44% decrease).

We analyzed spatial and temporal patterns in model consistency using the coefficient of variation (CV) between all 8 model predictions across Canada. For current predictions, we found that models predicted similar richness in the southern regions (southern Ontario and Quebec, etc.) but became less consistent towards northern regions (13% increase in CV from south to north, with the split defined by the 20% temperature isocline; Fig 4). Unsurprisingly, there was a trend of areas with lower sampling having higher CV (Fig 4 inset). Future projections followed a similar pattern where northern regions had more discrepancy (though southern Ontario also had high discrepancy in the future; Fig 4). Interestingly, overall discrepancy was consistently *lower* across Canada for future predictions than for current predictions (28% decrease; Fig 4).

**Figure 4.**
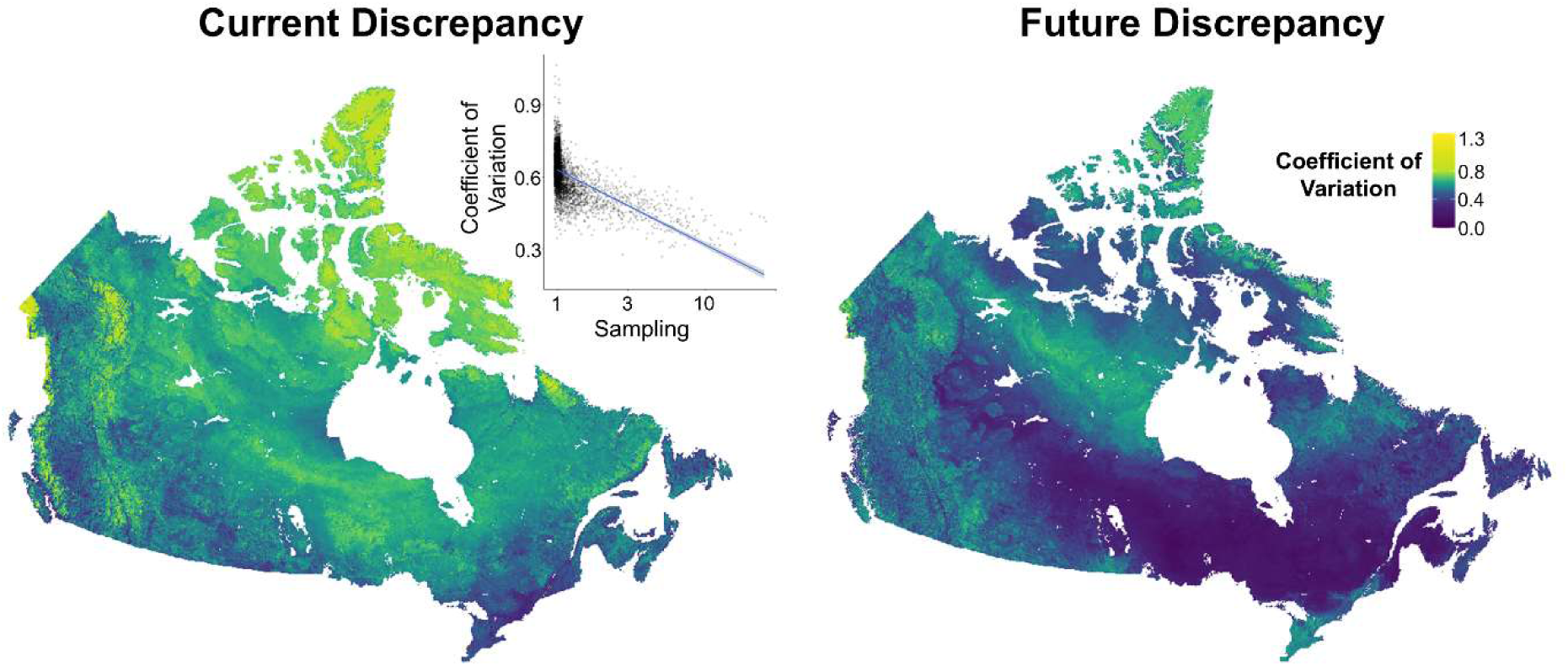
Spatial patterns of discrepancies between eight species distribution model methodologies for current and future (2080, RCP 8.5) terrestrial vertebrate richness estimates. Discrepancy is measured as the coefficient of variation among predictions (Variance/Mean). Inset plot shows relationship between sampling (number of GBIF occurrences +1; note that x-axis is log scale) and coefficient of variation (both metrics were aggregated to 50km grid cells to illustrate large scale relationship).

#### Range Shifts

Projections of range expansion were considerably less extreme after correction, by a factor of around 2. Species ranges in Canada were projected to increase in area by a factor of 3.48 on average (CI [3.23, 3.74]) with uncorrected models (NC). Species range expansion was less extreme with corrected models (TGB and BCC) increasing by a factor of 1.91 [1.77, 2.06] and 1.52 [1.43, 1.63] respectively.

Ranges can shift without changing in size, so we also report how much of the original range of a species remains stable in the future range. NC models predict that on average 69% [67%, 71%] of the original range will remain stable. TGB and BCC project 67% [66%, 69%] and 77% [75%, 78%] of the range remaining stable respectively.

To put this into perspective, we also compared average range sizes between models for the current scenarios. TGB estimated ranges that were twice as large (factor of 1.96 [1.87, 2.07]), and BCC five times as large (factor of 5.10 [4.84, 5.38]) as uncorrected models.

## Discussion

Overall, data limitation and bias along environmental gradients strongly influenced our ability to model present and future occurrence and richness for Canadian terrestrial vertebrates. As expected, bias-correction did generally improve prediction quality as evidenced by better validation against independent datasets. Nevertheless, different methods produced substantially different predictions of spatial biodiversity patterns and climate driven range shifts (high *prediction discrepancy*), with different models predicting a 2 to 5 fold difference in species richness in some cases. As expected, predictions in northern Canada exhibited more discrepancy than the south, but counter-intuitively, future projections exhibited less discrepancy than the present. In sum, our findings point to the importance of independent validation data and accounting for sampling patterns in environment space, especially when monitoring biodiversity in one of the Earth’s most rapidly warming regions (Rantanen *et al*. 2022).

### Multifaceted OCR validation

Our novel “occurrence-checklist-range” (OCR) validation confirmed that cross-validating with biased occurrences and background points can unfairly punish realistic models (there’s a high proportion of zeros in lesser sampled regions, even where bias-corrected models rightly predict high probabilities; Syfert *et al*. 2013). However, the magnitude of this effect in broad, real-world datasets has not been well demonstrated with explicit comparisons of cross-validations and independent validations. Additionally, models are still commonly evaluated using only cross-validation, since it is often seen as the only accessible data to compare against (Hijmans 2012). Our results suggest it is valuable and achievable to evaluate with any independent data that are available and have less spatial bias.

Using independent data at both fine (checklist) and coarse (range) scales is important because SDM predictions can be inaccurate in different ways and there are many ways to use SDM predictions in applications (Guillera-Arroita *et al*. 2015). Depending on the usage, different types of inaccuracies may be most detrimental to results, and a single metric might not capture this. For example, if SDMs are used to identify suitable habitat (i.e. to propose protected areas for an at-risk species), they should correctly discriminate between good and bad habitat within the range of the species (reflected by *checklist performance*). If SDMs are used to track range edges (i.e., to manage invasions or range-shifting species), they should predict low probabilities outside the true range (not necessarily captured by checklists but reflected by *range performance*). Beyond model performance, high *prediction discrepancy* may reveal certain shortcomings. Discrepancies in range shift metrics could indicate that area of occupancy trends are not reliable (i.e. for IUCN red list assessments; Guillera-Arroita *et al*. 2015), while discrepancies in future distributions could show they are unreliable (i.e. for designing resilient protected area networks; Santini *et al*. 2021). Quantifying multiple aspects of model performance, along with prediction discrepancy, gives us a more complete picture of the limitations and inaccuracies of SDM predictions in Canada.

### Inconsistent predictions in under-sampled environments

Our results build on past bias-correction studies by demonstrating the extent to which geographic sampling bias in GBIF data destabilizes biodiversity predictions in Canada. Although bias-correction methods improve *model performance*, they still failed to give consistent *predictions* of biodiversity patterns, changes in richness, and range shifts (high model uncertainty; Elith *et al*. 2002). This means that it’s difficult to know which, if any, of our methodologies produce trustworthy predictions. Selecting one method based on slightly better performance (in this case BCC) is not necessarily a good strategy as even BCC led to unexpected predictions (fairly even richness between south and north is not realistic; Raz *et al*. 2024). Without very clear evidence that one method should be trusted over the others, it’s best to acknowledge methodological uncertainty and assess when and where we have confidence in our predictions.

Prediction discrepancy was worse in poorly sampled environments (i.e., in the extensive north), indicating that even SDMs that perform well may be unreliable for large portions of the study area. This discrepancy coincides with regions expecting extreme warming (Bush & Lemmen 2019; Rantanen *et al*. 2022). This means that the very species which require effective climate change mitigation are also the ones for which anticipating changes (i.e. range contractions) will be most challenging (although some conservation planning efforts like spatial prioritization can be resilient to discrepancies in model predictions: Eckert *et al*. 2023; Guillera-Arroita *et al*. 2015).

Contrary to our expectations, discrepancy was *lower* for future projections. Although we expected that future unsampled combinations of environmental variables would force models to extrapolate more; (Elith *et al*. 2010a, Fitzpatrick & Hargrove 2009), this was superseded by the simple fact that future Canada will be warmer and we have better sampling coverage in warm environments (like the U.S.A.). Of course, future projections could still be inaccurate if the relationships between species and climate don’t hold into the future. Also, in other regions, unprecedented, unsampled climate conditions will still dominate and increase future discrepancy (Muhling *et al*. 2020). In Canada though, as climate change melts away the coldest, least sampled parts of environment space, we find a silver lining that SDM predictions may have *less* discrepancy. Here, sampling bias and climate change overlap *spatially* but not *temporally*.

These findings conflict somewhat with previous studies and highlight issues particular to this context of severe and large-scale sampling bias. Past studies focusing on Europe or the U.S.A. (and simulations) have some data bias but still have adequate data throughout most of environment space to guide model responses. Canada, however, is nearly devoid of northern data (on GBIF), so some findings differ in this understudied context. Firstly, some studies have found occurrence filtering to be a well-performing bias correction, and preferable to TGB for some contexts (Fourcade *et al*. 2014; Vollering *et al*. 2019). However, we found that gridded filtering alone (NC method) was inadequate for removing bias and performed poorly against independent validation data. Other studies have found that stronger bias-correction like TGB performs similarly to models without bias, or approximates true simulated distributions well (Baker *et al*. 2022; Barber *et al*. 2022; Inman *et al*. 2021). Bias-covariate correction has also been championed as a reliable method that circumvents problematic assumptions of TGB (Chauvier *et al*. 2021). Our results confirm that both methods improve predictions, but they also emphasize the discrepancies between methods. Although performance improves, predictions are still unreliable where models must extrapolate heavily for very poorly sampled parts of environment space (Meyer & Pebesma 2021). This situation is not unique to Canada, as many large areas of the globe exhibit a similar lack of data (Yesson *et al*. 2007). Our results indicate that global predictions based on GBIF data are likely to be unreliable in such regions, even with bias-correction.

Understanding and mitigating methodological uncertainty is crucial for realistically assessing climate-driven biodiversity change. Indicator metrics which rely on SDMs are increasingly being used to track conservation targets. For example, distribution estimates are required to calculate the species habitat index and species protection index indicators used in the Kunming-Montreal Global Biodiversity Framework (KMGBF. 2022, Griffith *et al*. 2024). To avoid mismanagement and retain trust in policy-relevant measurements of biodiversity, we must be realistic about model uncertainty (Cattarino *et al*. 2018). Going forward, we suggest three avenues for mitigating SDM uncertainty. First, sampling efforts should be targeted towards poorly sampled environments to optimize information gain (we suggest sampling schemes explicitly based on metrics of SDM uncertainty; Henrys *et al*. 2024). Second, we should improve the accessibility of data already collected in remote regions but is not published to open repositories (Camera traps, bioacoustics, collars, governmental and corporate environmental assessments, expert-based regional checklists, etc; King *et al*. 2012). Finally, we need to develop models that incorporate diverse data sources (each with different properties and assumptions) into coherent and better informed predictions in remote regions (Hui *et al*. 2024; Pollock *et al*. 2025), and share information between species using traits and biotic interactions to improve predictions for data-deficient species (Ovaskainen *et al*. 2024; Poggiato *et al*. 2025; Sharma *et al*. 2025). These steps will maximize the information that drives predictions in remote areas and reduce extrapolation. It’s clear from this analysis that in this northern context, bias correction is *necessary*, but *not sufficient* for making SDMs a reliable tool for measuring, predicting and mitigating climate impacts on species.

