## Supplemental Figures for "Where Climate Change and Sampling Bias Collide: Challenges of Predicting Biodiversity Change in Canada"

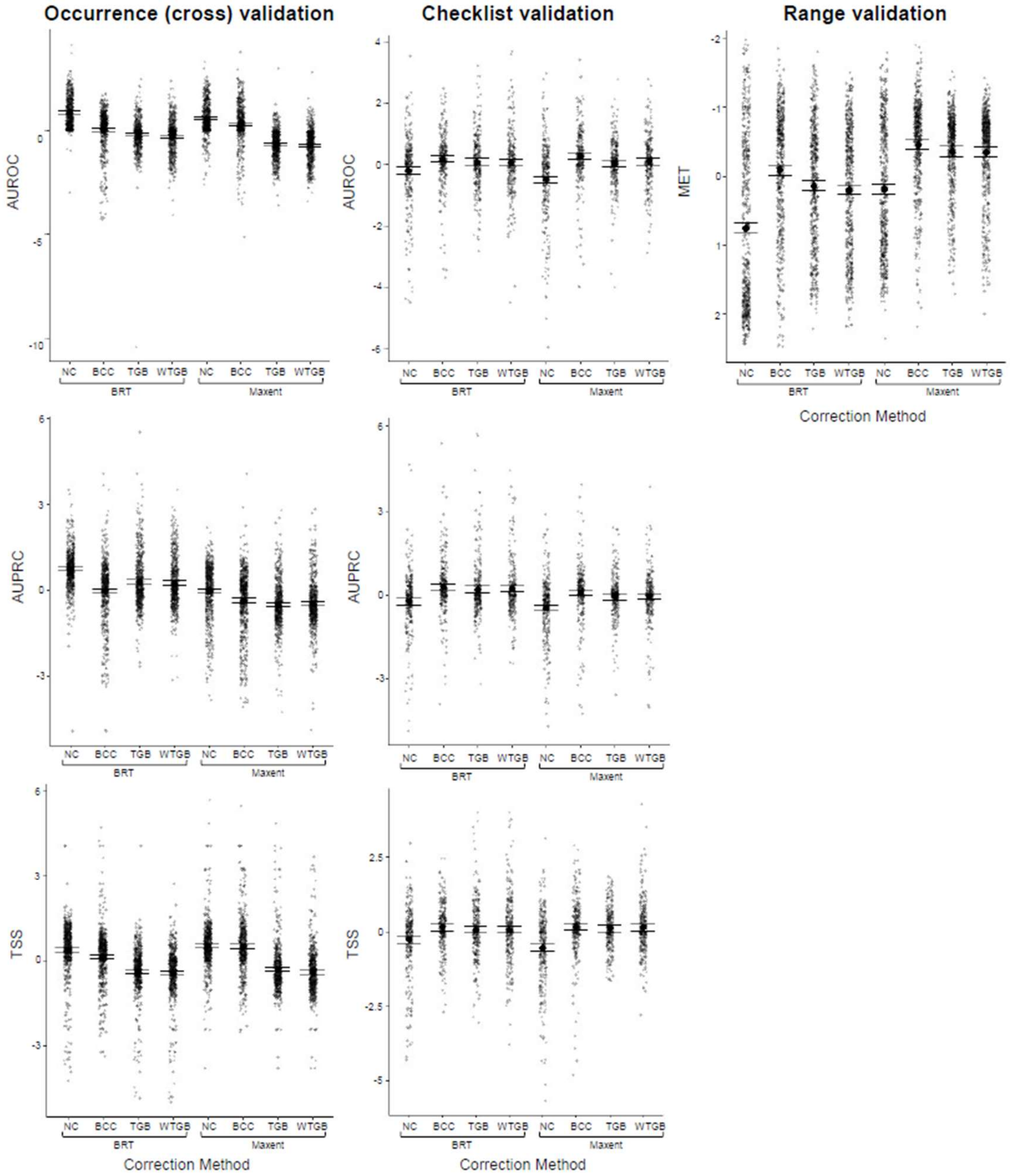


**Figure S1** Performance of different methodologies (bias correction methods: No Correction, Bias Covariate Correction, and Target Group Background, and Weighted TGB, algorithms: Boosted Regression Trees, Maxent) for 3 validation data sets; 30% hold-out sets from occurrences with psuedoabsences (cross-validation), presence/absence data from national park checklists, and range maps. Metrics for occurrence and checklist validations include Area Under the Receiver-Operator Curve, Area Under the Precision Recall Curve, and True Skill Statistic, (each centered within species and normalized across all species). Range validation uses the Minimum Energy Test (normalized within species). Large black dots represent the mean for each method, with 95% confidence interval as bars.

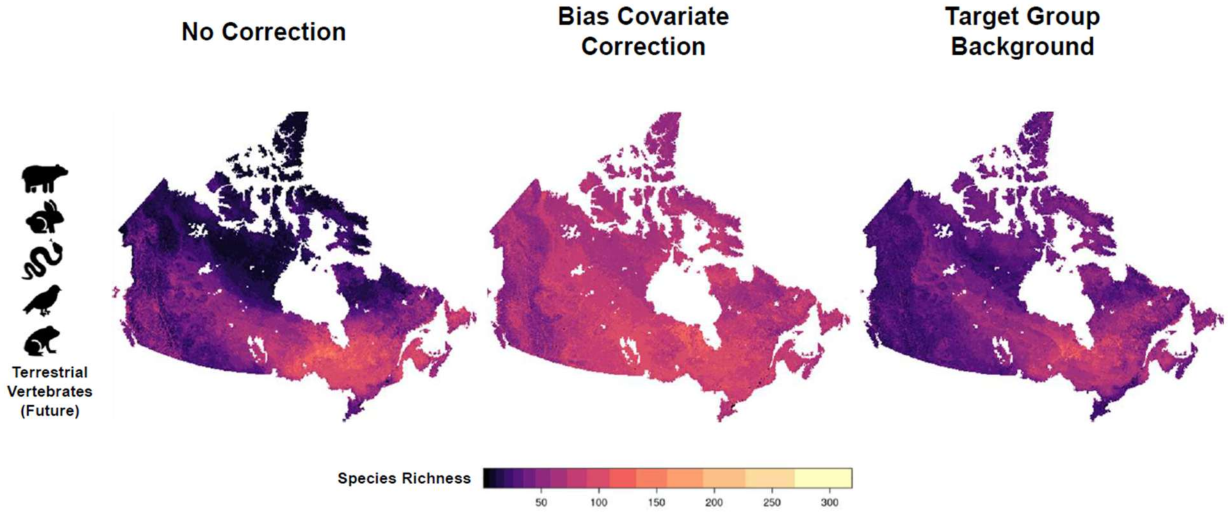


**Figure S2** Differences between three bias correction methods (all with boosted regression trees) of projected future (2080, RCP 8.5) species richness (as summed single species SDM predictions) of 698 terrestrial vertebrate species native to Canada.
